# Distinct Effects of the Male-Killing Bacteria *Wolbachia* and *Spiroplasma* and a Partiti-Like Virus in the Tea Pest Moth, *Homona magnanima*

**DOI:** 10.1101/2022.04.29.490121

**Authors:** Hiroshi Arai, Takumi Takamatsu, Shiou-Ruei Lin, Tetsuya Mizutani, Tsutomu Omatsu, Yukie Katayama, Madoka Nakai, Yasuhisa Kunimi, Maki N. Inoue

**Author notes:** Address correspondence to Hiroshi Arai,; Maki N. Inoue.

## Abstract

Male killing, the phenomenon of male death during development, is considered to be one of the advantageous strategies exerted by maternally transmitted microbes. Male killing has attracted interest in the fields of evolutionary biology and ecology for decades; however, little is known about its mechanism and origin. Here, we characterized and compared the effects of three distinct male killers, *Wolbachia* (Alphaproteobacteria), *Spiroplasma* (Mollicutes), and Osugoroshi virus (OGV) (Partitiviridae) in the tea pest moth *Homona magnanima* (Lepidoptera, Tortricidae). Regardless of the genetic sex (male: ZZ; female: ZW), female specific splice variants of the doublesex gene (*dsx*), a downstream regulator of the sex-determining gene cascade, was expressed in *H. magnanima* harbored either male-killing *Wolbachia* or *Spiroplasma*. However, OGV and non-male-killing *Wolbachia* did not alter *dsx* splicing. RNA sequencing and quantitative PCR assays demonstrated that male-killing *Wolbachia* impaired the host’s dosage compensation system by altering the global gene expression of the Z chromosome (corresponding to *Bombyx mori* chromosome 1 and 15) in males, whereas *Spiroplasma* did not affect dosage compensation. In contrast, the partiti-like virus OGVs did not affect sex-determination cascades or dosage compensation systems. Besides, male killers distinctly altered host gene expression and metabolomes associated with physiology, morphology, and diverse metabolic pathways. Moreover, *Wolbachia* and *Spiroplasma* infections triggered abnormal apoptosis only in male embryos. These findings suggest that distantly related microbes employ distinct machineries to kill identical host males, which have been acquired through independent evolutionary processes.

**Importance:** Male-killing caused by diverse microbes has attracted substantial attention. However, it remains unclear how such male killers have evolved similar phenotypes, in part because male-killing mechanisms have been studied using different insect models. Here, by comparing three phylogenetically distinct male killers, *Wolbachia, Spiroplasma*, and a partiti-like virus, in an identical host, we provide evidence that microbes can affect male viability through distinct machinery, demonstrating distinct evolutionary scenarios for microbes to acquire make-killing ability. These findings provide insight into new directions for studying microbe–host interactions.

## INTRODUCTION

Male killing (MK), a phenomenon of male death during development, is caused by microbes such as intracellular bacteria, microsporidia, and viruses in various insects (1, 2). The ecological and evolutionary significance of MK has attracted considerable attention for decades. Because male killers directly produce female-biased sex ratios in insects, the MK phenotype is considered to be a selfish strategy that promotes inheritance and propagation of microbes in nature (3–5). Moreover, male killers can have an overall advantage for the host insects if they benefit from the death of their male siblings through a reallocation of resources (3, 6). Indeed, several insects such as the ladybird beetle *Adalia bipunctata* have been shown to harbor phylogenetically distinct male killers (7–12).

A key question remains as to how phylogenetically distinct microbes achieved MK and evolved similar reproductive phenotypes in their host organisms. One proposed hypothesis is that microbes developed distinct MK mechanisms independently. Indeed, recent studies have indicated that MK could involve diverse mechanisms due to complex interactions between bacteria and host species (13–18). For instance, the MK bacterium *Wolbachia* (Alphaproteobacteria) in the fly *Drosophila bifasciata* causes male-specific abnormal apoptosis through a male dosage compensation system that adjusts the expression of sex-chromosome genes equally between males (XY) and females (XX) (15). In *Ostrinia* moths, *Wolbachia* inhibits male dosage compensation and defects sex-determination cascades by altering the transcription of the master regulator gene *doublesex (dsx*) (19–21). These findings suggest that even closely related microbes accomplish MK via distinct mechanisms in their hosts. In contrast, the bacterium *Spiroplasma poulsoni* (Mollicutes) causes abnormal apoptosis in *Drosophila melanogaster* males by utilizing the dosage compensation system, similar to the mechanism of *Wolbachia* in *D. bifasciata* (14, 22). Microbes sharing their niches frequently exchange their genes (23, 24). Thus, male killers coinfecting the same host may share similar MK mechanisms that have been acquired through interactions between host and microbes.

In this study, to clarify whether microbes employ distinct or similar MK machinery in an identical host, we compared the mechanisms of actions of three different male killers in the same host, the tea tortrix moth *Homona magnanima* (Tortricidae, Lepidoptera). Previous studies demonstrated that the partiti-like virus Osugoroshi virus (OGV) caused larval MK (25–27), but that *Spiroplasma* (28) and *Wolbachia* strain *w*Hm-t (29) caused embryonic MK in *H. magnanima*. In the present study, we identified that MK *Wolbachia* and *Spiroplasma* impaired sex-determination cascades and caused abnormal apoptosis in males. Unlike *Wolbachia*, *Spiroplasma* did not affect dosage compensation systems. Moreover, OGVs achieved MK without affecting either sex-determination or dosage compensation systems. Collectively, our findings highlight that distantly related microbes can exert MK via different mechanisms even in an identical host and have acquired MK abilities through independent evolutionary processes. We discuss the MK mechanisms and evolution of MK-inducing microbes in their host insects.

## RESULTS

### MK *Wolbachia* and *Spiroplasma*, but not OGVs, altered the splicing patterns of *dsx* in males

To evaluate the effects of the three male killers on sex determination in *H. magnanima*, we first determined the *dsx* transcript sequences. According to rapid-amplification of cDNA ends (RACE) assays, males had a variant *dsx-M* encoding the protein DSX-M, whereas females had eight splicing variants encoding three proteins, DSX-F1–3 (Fig. 1a). Male embryos (at 108 hours post-oviposition [hpo]) of the normal-sex ratio (NSR) line and the W^c^ line harboring non-MK *Wolbachia w*Hm-c (30) showed the *dsx*-M variant, whereas males infected with either *w*Hm-t (W^T12^, W^T24^, and W^TN10^) or *Spiroplasma* (S+) exhibited female-type *dsx (dsx-F*), encoding DSX-F1 or DSX-2 (Fig. 1b–c). In contrast, OGVs did not alter *dsx* splicing in male embryos (Fig. 1c) or in male larvae showing symptoms of viral infection (Fig. S1).

**Fig. 1.**
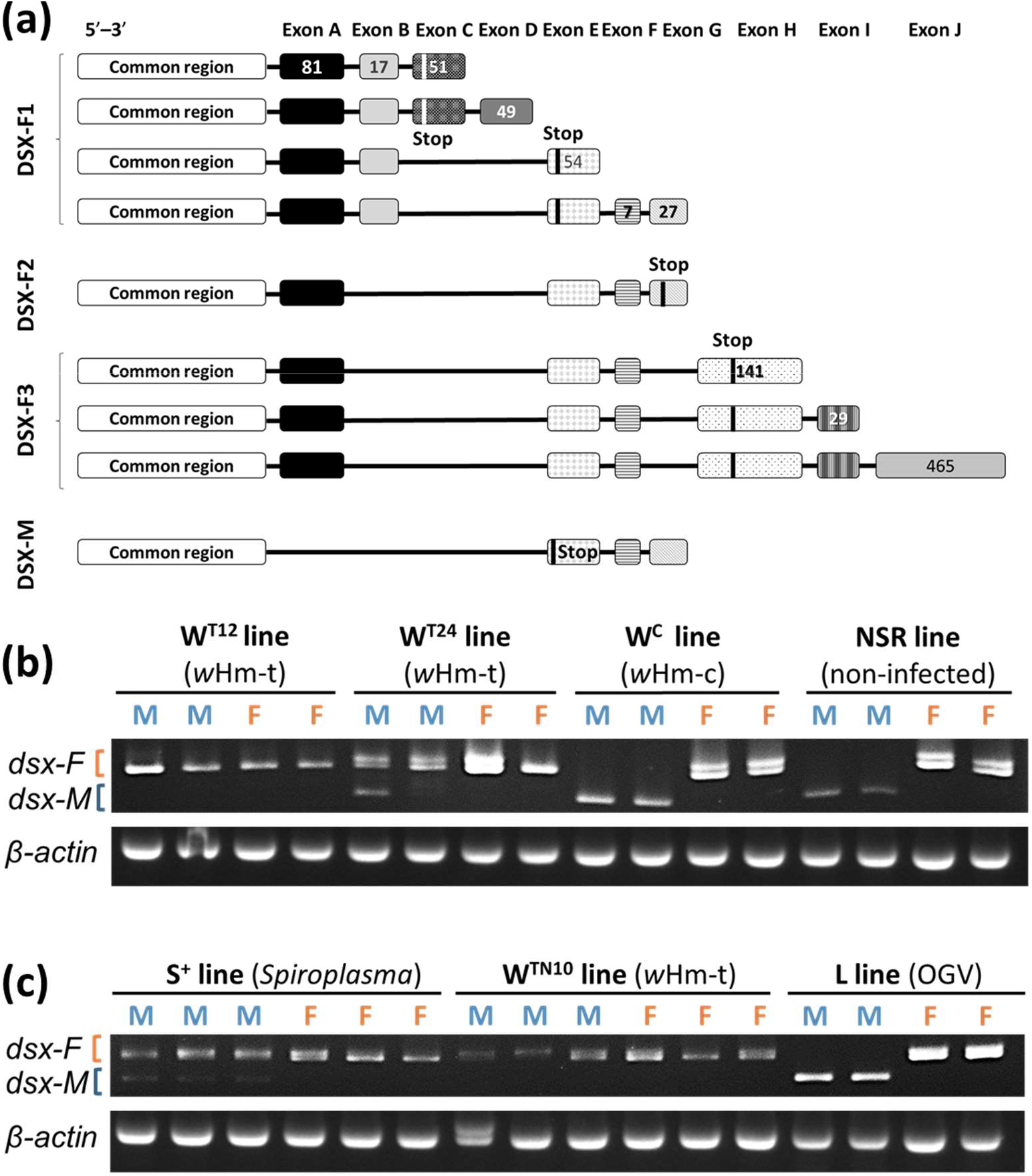
Effects of male killers on doublesex (*dsx*) mRNA splicing in *H. magnanima*. Nine *dsx* splicing variants in *H. magnanima*. Females had eight *dsx* variants encoding three DSX proteins (DSX-F1–3), and males showed a single variant (DSX-M). Rectangular boxes represent exons. The numbers inside the boxes indicate the numbers of base pairs. Stop codons in each variant are indicated with black or white bars. (b) *dsx*-splicing patterns in hosts harboring *w*Hm-t (W^T12^ and W^T24^) or *w*Hm-c (W^C^), and in Taiwan NSR lines. (c) *dsx*-splicing patterns in hosts harboring *Spiroplasma* (S^+^), *w*Hm-t (W^TN10^), or OGVs (L). The lengths of male-type *dsx (dsxM*, 377 bp) and female-type *dsx (dsxF*, 457 bp and 471 bp). β-actin was amplified as a control gene.

### Only MK *Wolbachia* impaired dosage compensation and expression of genes involved in sex determination systems

We evaluated dosage compensation effects in *H. magnanima* embryos harboring male killers using RNA-sequencing (RNA-seq) and quantitative polymerase chain reaction (qPCR) assays. A total of 54,071 contigs of the *de novo* assembled data (293,111 contigs, average length: 868.3 bp, total length: 254,508,797 bp) were annotated to *Bombyx mori* genes, which identified a transcriptional bias in putative Z chromosome genes (homologs of *B. mori* chromosome 1 [Ancestral Z chromosome: Anc-Z] and chromosome 15 [Neo Z chromosome: Neo-Z]) in *w*Hm-t-infected *H. magnanima* male embryos (Fig. 2a). In contrast, *Spiroplasma* and OGV infections did not show apparent fold-changes in the expression levels of Z-chromosome genes (Fig. 2b–c). The Z-linked triosephosphate isomerase (*HmTpi*) gene dose was 2-fold higher in males than in females, regardless of *Wolbachia* infection, confirming that males had homogametic sex chromosomes (ZZ), whereas females had heterogametic sex chromosomes (ZW) (Fig. 2f). Moreover, the Z chromosome genes *HmTpi* and kettin (*HmKettin*) were expressed at 2-fold higher levels in male embryos (108 hpo) harboring *w*Hm-t compared with those in females (Steel–Dwass test, *p* < 0.05). In contrast, *Spiroplasma* and OGVs infection did not affect the gene expression levels between the sexes (*p* > 0.05; Fig. 2e–f, Fig. S1).

**Fig. 2.**
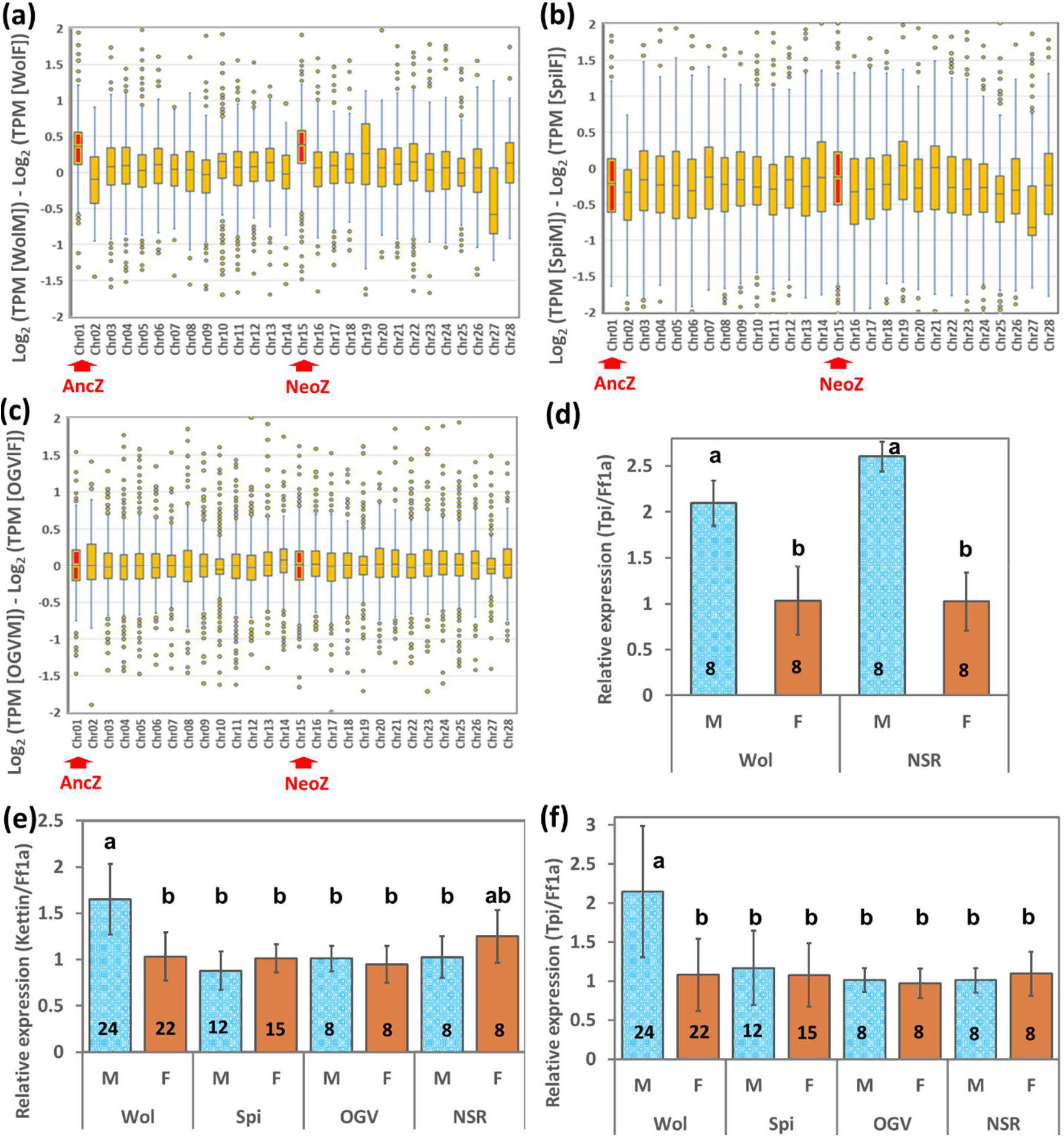
Three male killers affected *H. magnanima* dosage compensation differently. (a–c) Normalized expression levels (TPM) and chromosomal distributions of transcripts in *H. magnanima* embryos. RNA-seq data (108 hpo) were used to make the following comparisons: (a) W^12^ males versus W^T12^ females, (b) S^+^ males versus S^+^ females, and (c) L males versus L females. The chromosome number for each *H. magnanima* transcript-derived contig was assigned based on *B*. *mori* gene models. The boxes in the box-and-whisker diagrams represent the median and 25–75 percentile ranges of the expression ratios. (d–f) Quantification of Z-linked genes in *H. magnanima* embryos (108 hpo). The *Tpi* gene dose (d) was quantified to assess whether males have two Z chromosomes. *Kettin* (e) and *Tpi* (f) expression levels were normalized to that of the autosomal *Ef1α* gene. Error bars represent the standard error. The y axis indicates relative abundances adjusted to those of NSR females. Different letters indicate significant differences determined by the Steel–Dwass test (*p* < 0.05). The numbers inside the bars indicate replicates. *w*Hm-t: W^T12^ line; Spi: S^+^ line; OGV: L line; NSR: NSR line

To identify whether *w*Hm-t infection degrades the expression of genes involved in dosage compensation and sex-determination cascades in *B. mori* and *Ostrinia furnacalis* (21, 31), we compared time-dependent RNA-seq data of both *w*Hm-t-infected and uninfected *H. magnanima* embryos at 12, 36, 60, 84, and 108 hpo. Notably, *w*Hm-t infection degraded the expression of two masculinizer (*HmMasc*) isoforms, which showed sex-dependent expression in mature embryos (108 hpo) (Fig. 3a-b, S2). Although *w*Hm-t did not notably alter the expression dynamics of the P-element somatic inhibitor (*HmPSI*) gene, the expression of insulin-like growth factor II IGF-II mRNA binding protein (*HmIMP*) was upregulated by *w*Hm-t infection throughout embryogenesis (Fig. S2).

**Fig. 3.**
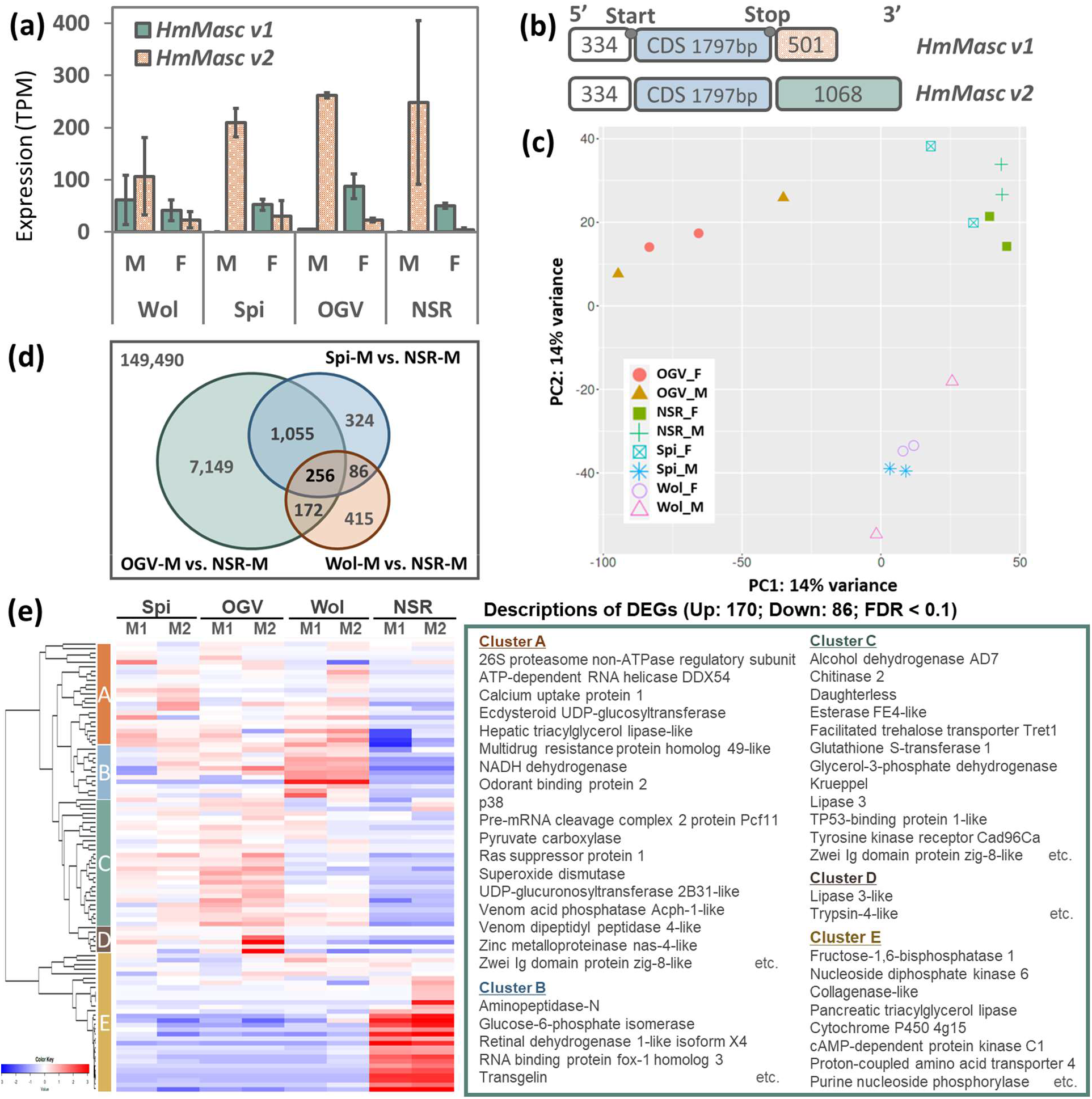
Gene expression patterns in *H. magnanima* harboring male killers. (a) Expression levels of *HmMasc* variants in male and female embryos (108 hpo). (b) Structures of *HmMasc* splice variants. The numbers inside the bars represent base pairs (bp). Both variants had identical 5’ untranslated regions (UTRs; white) and coding sequences (blue); however their 3’ UTRs (orange; green) were different. (c) Principal component analysis of host gene expression levels. Wol: W^T12^, Spi: Spi^+^, OGV: L, and NSR: NSR. (d) Numbers of deferentially expressed genes (DEGs) represented in Venn diagrams, with orange (W^T12^ males versus NSR males), blue (S^+^ males versus NSR males), and green (L males versus NSR males) shading. The numbers of non-DEGs are indicated outside the diagrams. (e) Classification of 256 DEGs shared by *H. magnanima* lines harboring male killers. The DEG-expression patterns are classified into four clusters with red (Up: upregulated) or blue (Down: downregulated) shading. The *H. magnanima* gene was assigned using *B*. *mori* gene models. The red and blue colors indicate upregulation and downregulation of DEGs, respectively, between W^T12^, S^+^, and L males versus NSR males.

### Male killers differently affected the host’s biological cascades

We identified 9457 differentially expressed genes (DEGs) altered by the three male killers in males by pairwise comparisons between eight groups of pooled embryos (uninfected, *w*Hm-t-infected, *Spiroplasma*-infected, and OGVs-infected males or females; 2–4 replicates; false-discovery rate [FDR] < 0.1) (Fig. 3c-d). The early male killers *w*Hm-t and *Spiroplasma* similarly altered the expression of genes, including enzymatic activities (e.g., protein kinases, aminopeptidase, ATPase) and detoxification genes (cytochrome P450), whereas OGVs affected genes involved in the metabolic pathways of males in different manners (Table S1, S2). Notably, *w*Hm-t upregulated a higher number of the host’s Z chromosome genes (accounting for 66 out of 336 DEGs, such as autophagy-protein 5 and multidrug resistance-associated protein) than those of males harboring either *Spiroplasma* (9 out of 391) or OGVs (35 out of 1040) (Table S2). *H. magnanima* harboring male killers shared 256 DEGs involved in stress responses (e.g., superoxide dismutase and p38), endocrine systems (e.g., Kruppel homolog 2), and signal responses (e.g., Ras suppressor protein, cAMP-dependent protein kinase) (Fig. 3c, Table S2).

To verify whether *Wolbachia* and *Spiroplasma* impair *H. magnanima* biosynthesis during embryogenesis, we quantified the host’s metabolome by capillary electrophoresis time-of-flight mass spectrometry (CE-TOF MS), which identified 273 metabolites. *Wolbachia* and *Spiroplasma* similarly upregulated dopamine-related substances (e.g., L-3,4-dihydroxyphenylalanine [DOPA] and dopamine) and sarcosine (a carcinogenesis-related substance) but downregulated 5-oxoproline and biopterin, which are involved in the nervous system. Of note, *w*Hm-t specifically upregulated the abundance of metabolites involved in sugar metabolism (e.g., *N*-acetylglucosamine) and the antioxidative stress response (e.g., *N^8^*-acetylspermidine; Table S3). In contrast, *Spiroplasma* specifically upregulated hypoxanthine (involved in the salvage pathway) and metabolites involved in energy synthesis (e.g., ATP, GTP, NADPH divalent, and cystine) but downregulated those involved in fat–amino acid metabolism (e.g., betaine).

### MK *Wolbachia* and *Spiroplasma*, but not OGVs, caused abnormal DNA damage during embryogenesis

Male embryos infected with *w*Hm-t or *Spiroplasma* (132 hpo) were fragile and exhibited nuclear condensation (Fig. 4a-b). We therefore further aimed to determine whether male killers underwent abnormal apoptosis during embryogenesis. Nuclear degradation occurred specifically in males (but not females) infected with either *w*Hm-t or *Spiroplasma* (Fig. 4c). The NSR line and L line harboring OGVs did not show apparent DNA fragmentation. The activities of caspase-3 (an apoptosis effector) were higher (Steel–Dwass test, *p* < 0.05) in *w*Hm-t-infected males (73539.9 ± 3625.8) and *Spiroplasma-infected* males (84434.8 ± 3773.8) than in non-infected males (48995 ± 3,773.8), *w*Hm-t-infected females (46697.9 ± 4357.6), and *Spiroplasma*-infected females (52128 ± 6536.4) (Fig. 4d). Moreover, terminal deoxynucleotidyl transferase (TdT) dUTP nick-end labeling (TUNEL) assays confirmed that males infected with *w*Hm-t or *Spiroplasma* exhibited abnormal nuclear segmentations (Fig. 4e, Fig. S3).

**Fig. 4.**
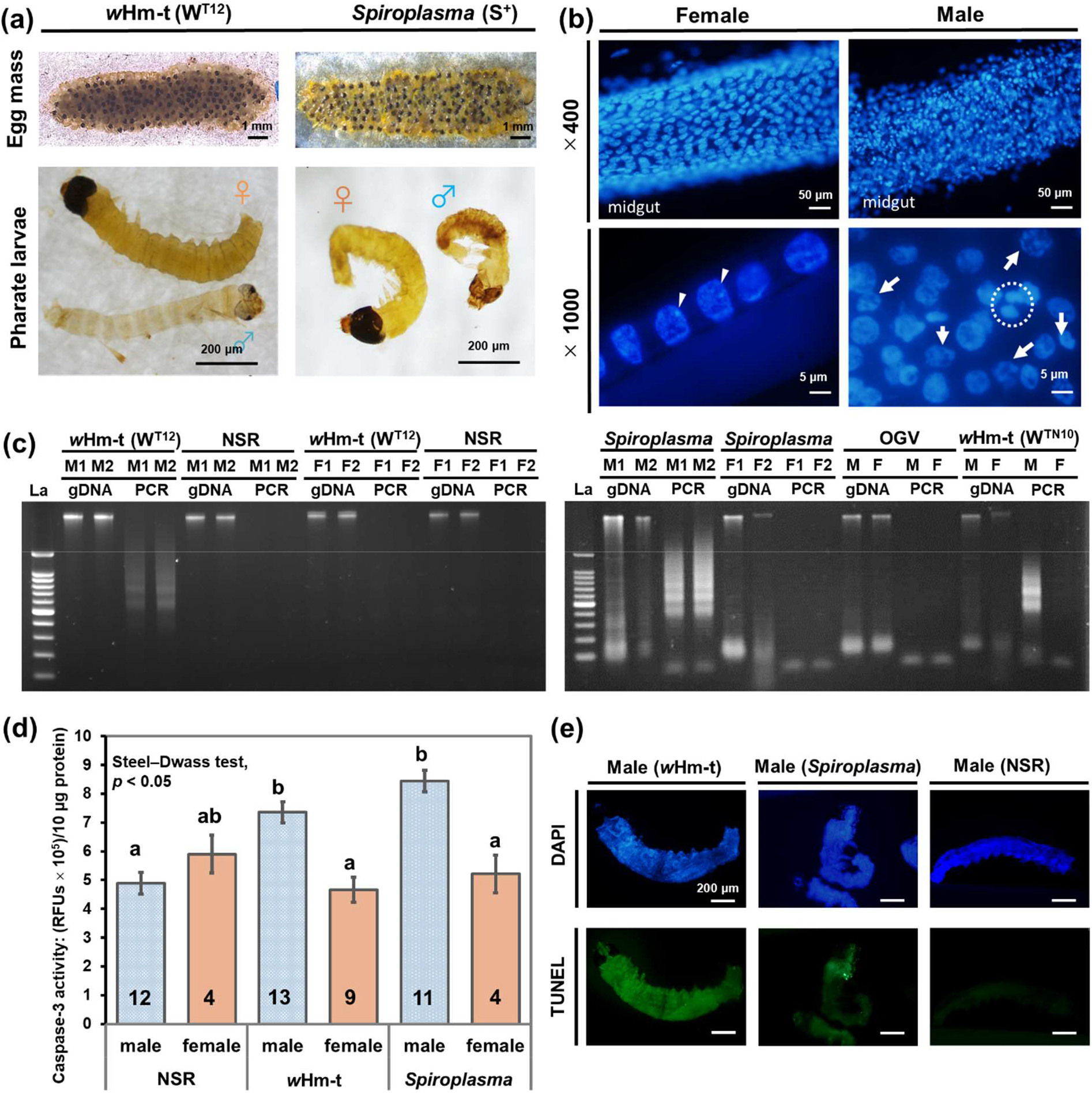
Male-killing *Wolbachia* and *Spiroplasma* caused abnormal apoptosis specifically in *H. magnanima* males. Morphological characteristics of embryos (132 hpo) extracted from W^T12^ and S^+^ egg masses. Male embryos were thinner and more fragile than the female embryo. (b) Representative image of the midgut of a W^T12^ embryo (132 hpo) stained with DAPI. Male embryos had abnormally shaped condensed nuclei (white arrows). The white arrowheads indicate heterochromatin (W chromosome). The broken white circle highlights the presumed chromatin bridge (x1000 magnification). (c) DNA ladders in the host lines. Genomic DNA (gDNA) and amplicons of laddered DNA (shown as “PCR”) from W^T12^, S^+^, L, and NSR embryos (132 hpo). La: 100-bp DNA ladder; M: male; F: female. (d) Caspase-3 activities in different host lines for NSR, W^T12^ (*w*Hm-t-positive) and S^+^ (*Spiroplasma*-positive) embryos (132 hpo). The numbers inside the bars indicate replicates. Different letters indicate significant differences between groups (Steel–Dwass test,*p* < 0.05). (e) TUNEL assays with whole-mounted NSR, W^T12^, and S^+^ male embryos (108 hpo). Green fluorescence indicates apoptosis and blue fluorescence indicates nuclei counterstained with DAPI. The sex of the embryos was confirmed by detecting the presence of absence of heterochromatin (W chromosome).

## DISCUSSION

### Do microbes require the sex-related machinery to kill insect males?

It has been hypothesized that microbes accomplish MK by targeting molecular mechanisms involved in sex determination and differentiation (4, 15, 32, 33). One key question is whether male killers have a unique mechanism that affects their host’s sex-related machinery. The dosage compensation system is often tightly associated with the sex-determination cascade in several insects (21, 31). *Wolbachia* in *O. furnacalis* disrupts dosage compensation and *dsx* splicing in males by downregulating the masculinizing gene *Masc* in an unknown manner (21). The present work in *H. magnanima* demonstrated that the early MK *Wolbachia* and *Spiroplasma* affected sex-determination cascades by altering *dsx* splicing, but only *Wolbachia* degraded the dosage compensation and the expression levels of two *Masc* variants in *H. magnanima* males. Therefore, our findings suggest that MK *Wolbachia* and *Spiroplasma* can alter *dsx* splicing in *H. magnanima* through different machinery (Fig. 5). We speculate that MK *Wolbachia* (i.e., *w*Hm-t and *w*Fur) generally alters *dsx* splicing by targeting *Masc* gene or its upstream factors in lepidopteran insects, while MK *Spiroplasma* alters *dsx* splicing by targeting downstream factors of *Masc* gene. Interestingly, OGVs did not affect dosage compensation or sex-determination cascades during embryogenesis or the larval stages, demonstrate that microbes do have distinct MK mechanisms even in the same host species.

**Fig. 5.**
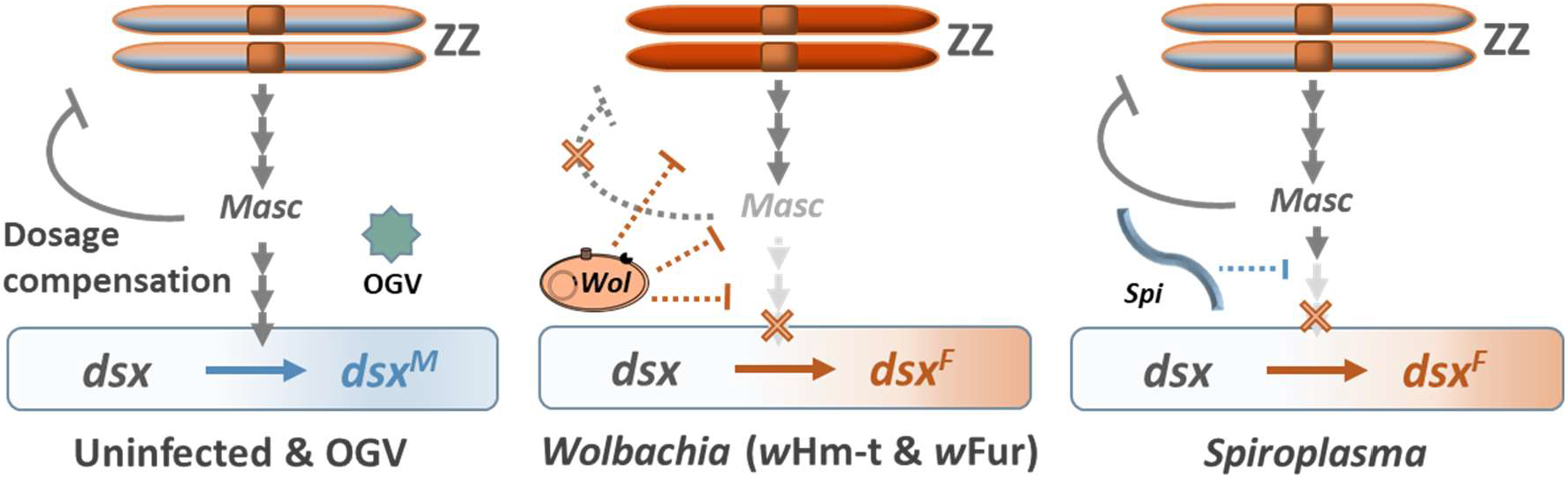
Proposed model for explaining the effects of male killers on the host’s sex-related machinery in Lepidoptera. The male-killing *Wolbachia* affected both sex-determination cascades and the dosage-compensation machinery, whereas *Spiroplasma* only affected sex determination in *H. magnanima*. *Wolbachia* possibly affects the dosage-compensation system and sex-determination cascade separately via different mechanisms or by targeting only the *Masc* gene and its upstream cascades, as predicted by Fukui et al. (21). In contrast, *Spiroplasma* utilizes a distinct, but unknown mechanism that directly affects the sex-determination cascade.

Improper dosage compensation has been thought to be the direct cause of *Wolbachia*-induced embryonic MK in *Ostrinia* moths (21). However, this assumption may require reconsideration as it is not entirely clear whether improper dosage compensation or feminization is the cause of MK. With this regard, our study demonstrated that MK *Spiroplasma* triggered alteration of *dsx* splicings even though dosage compensation was not affected. The irrelevance of dosage compensation effects in *Wolbachia* or *Spiroplasma*-induced embryonic MK is plausible because male mutants of the dosage compensation complex have been reported to die at late larval stages in *Drosophila* (34). Moreover, the adverse effects of improper dosage compensation may vary based on the gene composition of the Z chromosome. Although lepidopteran insects in general demonstrate synteny of the Z chromosomes, some lineages such as Tortricidae have large-scale Z chromosome rearrangements (35–40). Hence, improper dosage compensation induced by *Wolbachia* may result in different phenotypes depending on the gene composition of the Z chromosome in lepidopteran insects.

### How do male killers affect *H. magnanima* viability?

*Wolbachia* and *Spiroplasma* altered *dsx* splicing and triggered abnormal apoptosis in *H. magnanima* males. The *dsx* gene controls subsequent gene expression and sex-dependent characteristics in insects (41–43). As predicted by Kageyama and Traut (33), defects in the sex-determination cascades caused either by *Wolbachia* or *Spiroplasma* likely lead to mismatches between the genetic sex (ZZ: male) and phenotypic sex (*dsx* and subsequent gene-expression levels: female), which could affect the viability of males. Indeed, transcriptome and metabolome analyses have suggested that *Wolbachia* and *Spiroplasma* affect the motor functions (e.g., DOPA) (44), endocrine systems (e.g., juvenile hormone acid O-methyltransferase) (45), and antioxidative/antiaging activities (e.g., glutathione S-transferase and spermidine) (46) of males. Oxidative stress can cause DNA damage in various organisms and is associated with aging and dementia (47, 48). Such stresses caused by *Wolbachia* or *Spiroplasma* infection may induce DNA damage and lead to male embryo death in *H. magnanima*. Indeed, abnormal apoptosis is the cause of male-specific death in *Drosophila*, wherein *Spiroplasma* and *Wolbachia* trigger DNA damage to the male’s X chromosome (14, 15, 22).

In contrast to the early MK *Wolbachia* and *Spiroplasma*, OGVs killed larval or pupal males (24, 49) and did not affect male sex determination. Charlat et al. (50) suggested that the sex-determination cascade is not the sole target of MK in insects. The present work supports that prediction and indicates that OGVs may have a completely distinct MK mechanism without affecting the host’s sex-related machinery. Indeed, gene expression patterns in *H. magnanima* harboring OGVs differed from those in other host lines. OGVs may kill males by utilizing the differences in susceptibility between males and females toward viral factors (e.g., genes or viral loads), damaging male-specific organs (e.g., testes) or by impairing factors involved in sex dimorphism (downstream sex determination or maintenance systems such as hormone synthesis).

### Perspectives for the evolution of male killers and their host insect

In lepidopteran insects with a ZW/ZZ sex chromosome system, the W chromosome generally acts as an epistatic feminizer (51). In the present study, we highlighted that MK *Wolbachia w*Hm-t and *Spiroplasma* induce feminizing functions in *H. magnanima;* however, eliminating these microbes from the matrilines generates both males and females at the same ratio in the next generation (29), suggesting that the W chromosome of *H. magnanima* still acts as an epistatic feminizer. In contrast, eliminating MK *Wolbachia* generated “female-killing” in the next generation in *Ostrinia* moths (20, 52, 33, 52), due to the fact that the W chromosomes in *Wolbachia*-infected *Ostrinia* matrilines lose their female-determining function, which is compensated by MK (20, 33, 53). Similarly, *Wolbachia* is thought to be associated with feminization and the loss of W chromosomes in the butterfly *Eurema mandarina* (51). We speculate that, in Lepidoptera, the fundamental effect of MK *Wolbachia* is the disruption of sex determination cascades, which generates various reproductive phenotypes (i.e., MK and feminization) as a consequence of coevolutionary interactions between the host and microbes.

In summary, the present work reveals that phylogenetically distantly related microbes employ distinct MK mechanisms in an identical host *H. magnanima*. Our results suggest that microbes likely acquired MK abilities independently through different evolutionary processes. The specific factors responsible for MK (e.g., toxin genes) of each microbe should be examined in the future. Uncovering the complex interactions between insects and MK microbes will provide insights and new discoveries in host–parasite interactions, as well as evolutionary biology in the years to come.

## Materials and methods

### Insects

We previously established four *H. magnanima* lines (W^T12^, W^T24^, W^TN10^, and NSR) (29) collected from the Tea Research and Extension Station (Taoyuan-city, Taiwan) with permission from the Ministry of Agriculture, Forestry, and Fisheries (permission number 27–Yokohama Shokubou 891 and permission number 29–Yokohama Shokubou 1326) in 2015 and 2017. The W^T12^, W^T24^, and W^TN10^ lines are all-female matrilines harboring the MK *Wolbachia* (*w*Hm-t) strain. The NSR line exhibited a normal sex ratio (female male = 1:1) and was free of intracellular bacteria and OGVs (29). These Taiwanese *H. magnanima* lines were reared only at Tokyo University of Agriculture and Technology. The S^+^ and L lines are all-female and harbor *Spiroplasma* (28) and OGVs (27), respectively. The W^c^ line harbors a non-MK *w*Hm-c strain (30), which has a genotype identical to that of the MK strain *w*Hm-t (29). The insects were reared as described previously (30).

### Nucleic acid extractions from sex-determined insects

The genetic sexes of embryos (pharate larvae) were determined by observing W chromosomes (29, 33). Briefly, pharate larvae were dissected on glass slides with forceps, followed by fixation with methanol/acetic acid (50% v/v) and staining with lactic acetic orcein. The remaining tissues were stored in ISOGEN II (Nippon Gene) or cell lysis solution (10 mM Tris-HCl, 100 mM EDTA, and 1% SDS, pH 8.0) at – 80°C until subsequent extractions.

Total RNA and DNA were extracted from pharate larvae as described by Arai et al. (54). A total of 12 male or female pharate larvae were pooled and homogenized in the cell lysis solution or ISOGEN II. The DNA extraction procedure with cell lysis solution was performed as described by Arai et al. (30). To extract RNA, 600 μL of each homogenate was mixed with 240 μL of UltraPure Distilled Water (Invitrogen) and centrifuged at 12,000 ×*g* for 15 min at 4°C. Six hundred microliters of each supernatant was mixed with the same volume of isopropanol to precipitate the RNA, and then the resulting solutions were transferred to EconoSpin columns (Epoch Life Science) and centrifuged at 17,900 ×*g* for 2 min at 4°C. The RNAs captured in the column were washed twice with 80% ethanol and eluted in 20 μL of UltraPure Distilled Water (Invitrogen). Total RNA and DNA were also extracted from matured larvae and adults as described by Arai et al. (29). The extracted DNA and RNA were quantified using a Qubit v4.0 fluorometer (Invitrogen) and NanoPhotometer NP80 (Implen) and stored at −80°C until subsequent analysis.

### Sequencing and detection of the *H. magnanima dsx* gene

RNA samples extracted from female or male adults were reverse-transcribed via AMV Reverse Transcriptase XL (TaKaRa) using Oligo dT Adapter primers (Table S4). The resulting complementary DNA (cDNA) samples were amplified with the Hmdsx_long2F and Adapter primers using KOD -Plus-Ver.2 (Toyobo Co., Ltd.). The following PCR conditions were used: 2 min at 94°C, followed by 35 cycles of 10 s at 98°C, 30 s at 68°C, and 30 s at 72°C. The amplicons from the PCR were subjected to nested PCR with Hmdsx_long4F and adapter primers using Emerald Amp Max master mix under the following conditions: 2 min at 94°C, followed by 20 cycles of 10 s at 98°C, 30 s at 68°C, and 30 s at 72°C. The amplicons were sequenced as described by Arai et al. (29).

Detection of sex-specific *dsx* splicing variants was based on the procedure described by Sugimoto and Ishikawa (19). Briefly, total RNAs (100–300 ng) extracted from sex determined adults or pharate larvae (108 hpo) were reverse-transcribed using PrimeScript™ II Reverse Transcriptase (TaKaRa Bio Inc.) at 30°C for 10 min, 45°C for 60 min, and 70°C for 15 min. The cDNAs were then amplified using KOD-FX Neo (Toyobo Co., Ltd.) with the Hmdsx_long3F and Hmdsx_Mrev primer set (Table S4). The following PCR conditions were used: 94°C for 2 min, followed by 45 cycles of 98°C for 10 s and 68°C for 30 s. The amplicons were electrophoresed on 2.0% agarose Tris-borate-EDTA buffer (89 mM Tris-borate, 89 mM boric acid, 2 mM EDTA.) gels.

### RNA-seq analysis

We utilized 1.0 μg of total RNA extracted from W^T12^, S^+^, L, and NSR pharate larvae (108 hpo) or from W^T12^ and NSR egg masses (12, 36, 60, 84, and 108 hpo) to prepare mRNA-seq libraries. Two to four biological replicates were prepared for each treatment. First, mRNA was extracted from total RNA using the NEBNext Poly (A) mRNA Magnetic Isolation Module (New England Biolabs). The mRNA libraries were constructed with NEBNext Ultra II RNA Library Prep Kit for Illumina (New England Biolabs) following the manufacturer’s protocol to prepare 300-bp RNA fragments for 150 bp paired-end (PE150) analysis. The generated sequence data from each library were trimmed by eliminating (1) adaptor sequences, (2) reads harboring non-determined sequences exceeding 10%, and (3) sequences harboring low-quality nucleotides (Qscore < 5) spanning >50% of the read length by Novogen (Beijing, China). The trimmed reads were subjected to subsequent analyses.

### *De novo* assembly and DEG analysis

The trimmed reads were assembled *de novo* to generate a transcriptome database for *H. magnanima* using Trinity (55) and NAAC Galaxy with the default parameters. The expression levels of contigs were analyzed using Kallisto software (56) and iDep9 (http://bioinformatics.sdstate.edu/idep/). The DEGs of *H. magnanima* identified under different conditions were annotated using blast2go (57) and *B. mori* protein datasets (58), which were downloaded from KAIKObase (https://kaikobase.dna.affrc.go.jp). The annotated Gene Ontology terms were compared to identify biological processes that were enriched after infections by male killers by gene set-enrichment analysis (GSEA).

### Analyzing dosage compensation and quantifying genes on the Z chromosome

The effects of male killers on dosage compensation were verified by measuring gene expression differences, as described by Fukui et al. (21) and Gu et al. (59). To assess fold-changes in gene expression levels, transcripts per million (TPM) values were calculated using Kallisto software for all annotated *B. mori* gene homologs (assembled *H. magnanima* contigs) in both sexes. The binary logarithms of the TPM differences between males and females on *B. mori* chromosomes 1 to 28 were plotted.

Z chromosomal gene expression in *H. magnanima* was further quantified by reverse transcription-qPCR. Homologs of the *B. mori* Z chromosomal genes *Kettin* and *Tpi* were extracted from *H. magnanima de novo*-assembled data by performing a BLASTx search. The primer sequences used to quantify these genes are shown in Table S4. The gene doses were quantified using DNA extracted from males and females. Next, 100 ng RNA extracted from 12 male or female pharate larvae was reverse-transcribed using PrimeScript II reverse transcriptase (TaKaRa) at 50°C for 30 min, followed by denaturation at 95°C for 5 min. The cDNA was used to quantify relative gene expression levels, with normalization to the control gene elongation factor 1a (*ef1a;* Table S4). The mean cycle threshold (Ct) values of dual samples were calculated for at least eight replicates, and both ΔCt (CtAve Z gene – CtAve ef1a) and ΔΔCt (ΔCtmale – ΔCtfem) values were calculated. The dosage (number of Z chromosomes) was estimated based on the 2^−ΔΔCt^ method, as described by Sugimoto et al. (20).

### Quantification of genes involved in sex-determination cascades

*H. magnanima IMP*, *PSI*, and *Masc* gene homologs associated with the sex-determination system in *B. mori* were annotated using BLASTx (bit-score > 200) with protein datasets obtained from KAIKObase. The time-dependent gene expression levels in males and females during embryogenesis up to pharate larvae (12–108 hpo) were plotted using the TPM values, as described above.

### Metabolome analysis

Twenty milligram egg masses (108 hpo) from the NSR, W^T12^, and S^+^ lines were homogenized separately in 50% acetonitrile and centrifuged at 4°C and 2300 x*g* for 5 min. Then, 400 μL of each supernatant was filtered through ultrafiltration tubes (Ultrafree MC PLHCC, 5 kDa, Human Metabolome Technologies) at 4°C and 9100 x*g* for 120 min. The filtrates were resuspended in 100 μL Milli-Q water and analyzed using an Agilent CE-TOFMS system (Agilent Technologies, CA, USA). The relative peak area (sample quantity × peak area/area of the internal standard) was converted using MasterHands software ver.2.17.1.11, and peaks with a signal-to-noise ratio of >3 were detected from the CE-TOF MS data. Based on the mass-to-charge ratio (*m/z*) and migration time, each peak was annotated using Human Metabolome Technologies metabolome libraries.

### Observing apoptosis in mature male and female embryos

DNA segmentations were visualized as described by Staley et al. (60) using DNA extracted from W^T12^, S^+^, L, and NSR pharate larvae (132 hpo). Briefly, 100 ng DNA, 1 μL 24-bp linker (1 nM), 1 μL 12-bp linker (1 nM) (Table S4), and 5 μL 2x buffer for T4 DNA Ligase (Promega) were mixed and incubated at 55°C for 10 min, cooled down gently to 10°C for 55 min, and then incubated at 10°C for 10 min. The reaction mixture was incubated with 1 μL T4 DNA ligase (3 U/μL) for 15 min at 25°C. Then, 1.5 μL of ligated reactant (15 ng DNA) was amplified using TaKaRa Ex-Taq (TaKaRa) with the following conditions: 72°C for 5 min and 25 cycles of 94°C for 1 min and 72°C for 3 min. The genomic DNA and PCR amplicons were electrophoresed on 1.5% TBE agarose gels to visualize the ladders.

To quantify the caspase-3 activity, W^T12^, S^+^, and NSR pharate larvae (132 hpo) were first homogenized in 120 –L of 1x lysis buffer (10 mM Tris-HCl, pH 7.5, 100 mM NaCl, 1 mM EDTA, 0.01% Triton™ X-100). After centrifugation at 3000 x*g* for 5 min, 100 μL of each supernatant or 1x lysis buffer was used as a sample or background control for the following assays, respectively. The caspase-3 activity was quantified using EnzChek™ Caspase-3 Assay Kit #1 (Invitrogen) following the manufacturer’s protocol. The fluorescence intensity (excitation/emission at ~342/441 nm, respectively) was quantified using 1420 ARVOTM MX-fla Multilabel Counter (Perkin Elmer).

To visualize DNA damage by performing TUNEL assays, mature embryos (108 hpo) and midguts of pharate larvae (132 hpo) were fixed with 4% (w/v) paraformaldehyde phosphate-buffered saline with Tween-20 (PBST; 137 mM NaCl, 8.1 mM Na_2_HPO_4_, 2.68 KCl, 1.47 KH_2_PO_4_, 0.05% Tween-20, pH 7.4 as described in Arai et al. (61) solution for 15 min, washed twice with PBST for 5 min, incubated with 100 μL proteinase K solution (20 μg/mL) for 20 min, and washed twice with PBST for 5 min. Positive controls were treated with 50 μL DNase solution (TaKaRa) for 20 min. The TUNEL assays were performed using the DeadEnd™ Fluorometric TUNEL System (Promega) following the manufacturer’s protocol. The TUNEL-stained samples were immersed in 4’,6-diamidino-2-phenylindole (DAPI) solution (1 μg/mL; Dojindo) for 5 min and then washed twice with PBST for counterstaining. Samples were observed by fluorescence microscopy with ProLong™ Diamond Antifade Mountant. Sexes were determined based on the presence of the W chromosome.

## Data analysis and accessibility

Z chromosomal gene expression levels and caspase activities were analyzed using the Steel–Dwass test in JMP software v9 (SAS, Cary, NC, USA). The *Masc, Imp, PSI*, and *dsx* sequences of *H. magnanima* were deposited in GenBank under accession numbers LC701633 to LC701646. High-throughput sequencing data are available under accessions DRA013555 and PRJDB13118 (BioProject). Assembled data are accessible under PRJDB13118_Homona_TSA (ICSI01000001-ICSI01293111).

## Acknowledgements

We thank Dr. Daisuke Kageyama (The National Agriculture and Food Research Organization, NARO) and Dr. Toshiyuki Harumoto (The HAKUBI Center for Advanced Research, Kyoto University) for their kind advice regarding our research.

The authors wish to acknowledge support from the Japan Society for the Promotion of Science (JSPS) Research Fellowships for Young Scientists [Grant Number 19J13123 and 21J00895] and the Leading Program Research Foundations from the Ministry of Education, Culture, Sports, Science and Technology/Tokyo University of Agriculture and Technology to Hiroshi Arai.

## Data Accessibility and Benefit-Sharing

### Data Accessibility

Sequence data are deposited in the DRA013555 and PRJDB13118 (BioProject).

### Benefits Generated

The *H. magnanima* were collected from tea plantations at Tea Research and Extension Station (Taoyuan City, Taiwan) and were imported with permission from the Ministry of Agriculture, Forestry and Fisheries (No. 27 - Yokohama Shokubou 891 and No. 297 - Yokohama Shokubou 1326). Taiwanese *H. magnanima* were maintained only at Tokyo University of Agriculture and Technology. A research collaboration was developed with scientists from the countries providing genetic samples, all collaborators are included as co-authors, the results of research have been shared with the provider communities and the broader scientific community. More broadly, our group is committed to international scientific partnerships, as well as institutional capacity building.

## Author Contributions

In this work, HA had responsibility for the decision to submit for publication, wrote the paper, designed and conducted most experiments, and analyzed the data. SRL contributed to the research designation and discussion. TT conducted a part of the RACE experiments. TM, TO, and YK helped with and advised on the design of the RNA-seq experiments. MN and YK supported all experiments and contributed to the discussions related to the entire study. MNI contributed to the entire organization of this study and had responsibility for the decision to submit for publication.

## Legends of Supplemental Materials

**Fig. S1 The OGVs did not affect the host’s sex-related machinery during larval development.**

(a–d) Morphologies of last-instar larvae of *H. magnanima*. The males harboring OGVs (a) showed carcinoma-like tissue, which was not observed in females of the same line (b), NSR males (c), or NSR females (d). The blue arrows indicate testes, and the orange arrowheads indicate carcinoma-like tissues. (f) Expression levels of the Z-linked genes *Tpi* and *Kettin*, normalized to expression of the autosomal *Eflα* gene in the last-instar larvae. The numbers inside the bars indicate replicates. (g) The splicing patterns of the *dsx* gene in NSR (OGV−) and L (OGV+) lines. M: males; F: females

**Fig. S2. Effects of MK *Wolbachia* on genes involved in the host’s sex determination**

(a) Model of the sex-determination cascade in *B. mori* based on a previous report by Kiuchi et al. (2014). The feminizing PIWI-interacting RNA, *Fem*, interfered with *Masc* functions in females. *Masc* facilitates expression of the male splicing gene *BmIMP*, and both *BmIMP* and *BmPSI* determine *dsx* splicing in males. Z: Z chromosome; W: W chromosome. (b–e) Expression dynamics of *HmPSI* (b), *HmIMP* (c), and *HmMasc* (d-e) in the W^T12^ and NSR lines. The expression levels were quantified every 24 h from 12 h post-oviposition (hpo) to 108 hpo.

**Fig. S3 Apoptosis signals in matured male embryos**

TUNEL assays were performed in NSR and W^T12^ tissues (108 hpo males and females) to visualize apoptosis (green). DAPI was used as the counterstain (blue). The sex of each whole-mounted embryo was confirmed after TUNEL staining based on W chromosome observations.

**Table S1. Enriched pathways associated with differentially expressed genes (DEGs) by Gene ontology (GO) analysis in *H. magnanima***

**Table S2. Differentially expressed host genes in *Wolbachia* infected males.**

**Table S3. Metabolites showing >2-fold changes between the W^T12^ and NSR or S+ and NSR lines.**

**Table S4. Sequences and related information of the primers used in this study.**

## References

1. Kageyama D, Narita S, Watanabe M. 2012. Insect sex determination manipulated by their endosymbionts: incidences, mechanisms and implications. Insects 3:161–199.

2. Ma WJ, Vavre F, Beukeboom LW. 2014. Manipulation of arthropod sex determination by endosymbionts: diversity and molecular mechanisms. Sex Dev 8:59–73.

3. Hurst LD. 1991. The incidences and evolution of cytoplasmic male killers. Proc Royal Soc B 244:91–99.

4. Hurst GD, Jiggins FM. 2000. Male-killing bacteria in insects: mechanisms, incidence, and implications. Emerg Infect Dis 6:329–336.

5. Hornett EA, Charlat S, Duplouy AMR, Davies N, Roderick GK, Wedell N, Hurst GDD. 2006. Evolution of male-killer suppression in a natural population. PLoS Biol 4:e283.

6. Werren JH, Baldo L, Clark ME. 2008. *Wolbachia*: master manipulators of invertebrate biology. Nat Rev Microbiol 6:741–751.

7. Werren JH, Hurst GDD, Zhang W, Breeuwer R, Stouthamer R, Majerus ME. 1994. Rickettsial relative associated with male killing in the ladybird beetle (*Adalia bipunctata*). J Bacteriol 176:388–394.

8. Hurst GDD, Jiggins FM, von der Schulenburg JHG, Bertrand D, West SA, Goriacheva II, Zakharov IA, Werren JH, Stouthamer R, Majerus MEN. 1999. Male–killing *Wolbachia* in two species of insect. Proc Royal Soc B 266:735–740.

9. Hurst GDD, von der Schulenburg JHG, Majerus TM, Bertrand D, Zakharov IA, Baungaard J, Völkl W, Stouthamer R, Majerus ME. 1999. Invasion of one insect species, *Adalia bipunctata*, by two different male-killing bacteria. Insect Mol Biol 8:133–139.

10. Kageyama D, Nishimura G, Ohno S, Takanashi T, Hoshizaki S, Ishikawa Y. 2004. *Wolbachia* infection and an all-female trait in *Ostrinia orientalis* and *Ostrinia zaguliaevi*. Entomol Exp Appl 111:79–83.

11. Tabata J, Hattori Y, Sakamoto H, Yukuhiro F, Fujii T, Kugimiya S, Mochizuki A, Ishikawa Y, Kageyama D. 2011. Male killing and incomplete inheritance of a novel *Spiroplasma* in the moth *Ostrinia zaguliaevi*. Microb Ecol 61:254–263.

12. Takamatsu T, Arai H, Abe N, Nakai M, Kunimi Y, Inoue MN. 2021. Coexistence of two male-killers and their impact on the development of oriental tea tortrix *Homona magnanima*. Microb Ecol 81:193–202.

13. Harumoto T, Anbutsu H, Lemaitre B, Fukatsu T. 2016. Male-killing symbiont damages host’s dosage-compensated sex chromosome to induce embryonic apoptosis. Nat Commun 7:1–12.

14. Harumoto T, Lemaitre B. 2018. Male-killing toxin in a bacterial symbiont of *Drosophila*. Nature 557:252–255.

15. Harumoto T, Fukatsu T, Lemaitre B. 2018. Common and unique strategies of male killing evolved in two distinct *Drosophila* symbionts. Proc Royal Soc B 285:20172167.

16. Reynolds LA, Hornett EA, Jiggins CD, Hurst GDD. 2019. Suppression of *Wolbachia*-mediated male-killing in the butterfly *Hypolimnas bolina* involves a single genomic region. PeerJ 7:e7677.

17. Perlmutter JI, Bordenstein SR, Unckless RL, LePage DP, Metcalf JA, Hill T, Martinez J, Jiggins FM, Bordenstein SR. 2019. The phage gene *wmk* is a candidate for male killing by a bacterial endosymbiont. PLoS Pathog 15:e1007936.

18. Ferree PM, Avery A, Azpurua J, Wilkes T, Werren JH. 2008. A bacterium targets maternally inherited centrosomes to kill males in *Nasonia*. Curr Biol 18:1409–1414.

19. Sugimoto TN, Ishikawa Y. 2012. A male-killing *Wolbachia* carries a feminizing factor and is associated with degradation of the sex-determining system of its host. Biol Lett 8:412–415.

20. Sugimoto TN, Kayukawa T, Shinoda T, Ishikawa Y, Tsuchida T. 2015. Misdirection of dosage compensation underlies bidirectional sex-specific death in *Wolbachia-infected Ostrinia scapulalis*. Insect Biochem Mol Biol 66:72–76.

21. Fukui T, Kawamoto M, Shoji K, Kiuchi T, Sugano S, Shimada T, Suzuki Y, Katsuma S. 2015. The endosymbiotic bacterium *Wolbachia* selectively kills male hosts by targeting the masculinizing gene. PLoS Pathog 11:e1005048.

22. Veneti Z, Bentley JK, Koana T, Braig HR, Hurst GDD. 2005. A functional dosage compensation complex required for male killing in *Drosophila*. Science 307:1461–1463.

23. Wiedenbeck J, Cohan FM. 2011. Origins of bacterial diversity through horizontal genetic transfer and adaptation to new ecological niches. FEMS Microbiol Rev 35:957–976.

24. Kent BN, Bordenstein SR. 2010. Phage WO of *Wolbachia:* lambda of the endosymbiont world. Trends Microbiol 18:173–181.

25. Morimoto S, Nakai M, Ono, A, Kunimi Y. 2001. Late male-killing phenomenon found in a Japanese population of the oriental tea tortrix, *Homona magnanima*(Lepidoptera: Tortricidae). Heredity 87:435–440.

26. Nakanishi K, Hoshino M, Nakai M, Kunimi Y. 2008. Novel RNA sequences associated with late male killing in *Homona magnanima*. Proc Royal Soc B 275:1249–1254.

27. Fujita R, Inoue MN, Takamatsu T, Arai H, Nishino M, Abe N, Itokawa K, Nakai M, Urayama S, Chiba Y, Amoa-Bosompem M, Kunimi Y. 2021. Late male-killing viruses in *Homona magnanima* identified as Osugoroshi Viruses, novel members of *Partitiviridae*. Front Microbiol 11:620623.

28. Tsugeno Y, Koyama H, Takamatsu T, Nakai M, Kunimi Y, Inoue MN. 2017. Identification of an early male-killing agent in the oriental tea tortrix, *Homona magnanima*. J Hered 108:553–560.

29. Arai H, Lin SR, Nakai M, Kunimi Y, Inoue MN. 2020. Closely related male-killing and nonmale-killing *Wolbachia* strains in the oriental tea tortrix *Homona magnanima*. Microb Ecol 79:1011–1020.

30. Arai, H, Hirano T, Akizuki N, Abe A, Nakai M, Kunimi Y, Inoue MN. 2019. Multiple infection and reproductive manipulations of *Wolbachia* in *Homona magnanima* (Lepidoptera: Tortricidae). Microb Ecol 77:257–266.

31. Kiuchi T, Koga H, Kawamoto M, Shoji K, Sakai H, Arai Y, Ishihara G, Kawaoka S, Sugano S, Shimada T, Suzuki Y, Suzuki MG, Katsuma S. 2014. A single female-specific piRNA is the primary determiner of sex in the silkworm. Nature 509:633–636.

32. Anbutsu H, Fukatsu T. 2003. Population dynamics of male-killing and non-male-killing spiroplasmas in *Drosophila melanogaste*r. Appl Environ Microbiol 69:1428–1434.

33. Kageyama D, Traut W. 2004. Opposite sex–specific effects of *Wolbachia* and interference with the sex determination of its host *Ostrinia scapulalis*. Proc Royal Soc B 271:251–258.

34. Baker BS, Gorman M, Marín I. 1994. Dosage compensation in Drosophila. Annu Rev Genet 28:491–521.

35. Erickson JW, Quintero JJ. 2007. Indirect effects of ploidy suggest X chromosome dose, not the X:A ratio, signals sex in *Drosophila*. PLoS Biol 5:e332.

36. Lee J, Kiuchi T, Kawamoto M, Shimada T, Katsuma S. 2015. Identification and functional analysis of a Masculinizer orthologue in *Trilocha varians* (Lepidoptera: Bombycidae). Insect Mol Biol 24:561–569.

37. Nguyen P, Sýkorová M, Šíchová J, Kůta V, Dalíková M, Frydrychová RČ, Neven LG, Sahara K, Marec F. 2013. Neo-sex chromosomes and adaptive potential in tortricid pests. Proc Natl Acad Sci USA 110:6931–6936.

38. Fraïsse C, Picard MAL, Vicoso, B. 2017. The deep conservation of the Lepidoptera Z chromosome suggests a non-canonical origin of the W. Nature Commun 8:1–9.

39. Wan F, Yin C, Tang R, Chen M, Wu Q, Huang C, Qian W, Rota-Stabelli O, Yang N, Wang S, Wang G, Zhang G, Guo J, Gu LA, Chen L, Xing L, Xi Y, Liu F, Lin K, Guo M, Liu W, He K, Tian R, Jacquin-Joly E, Franck P, Siegwart M, Ometto L, Anfora G, Blaxter M, Meslin C, Nguyen P, Dalíková M, Marec F, Olivares J, Maugin S, Shen J, Liu J, Guo J, Luo J, Liu B, Fan W, Feng L, Zhao X, Peng X, Wang K, Liu L, Zhan H, Liu W, Shi G, Jiang C, Jin J, Xian X, Lu S, Ye M, Li M, Yang M, Xiong R, Walters JR, Li F. 2019. A chromosome-level genome assembly of *Cydia pomonella* provides insights into chemical ecology and insecticide resistance. Nature Commun 10:1–14.

40. Uchibori-Asano M, Jouraku A, Uchiyama T, Yokoi K, Akiduki G, Suetsugu Y, Kobayashi T, Ozawa A, Minami S, Ishizuka C, Nakagawa Y, Daimon T, Shinoda T. 2019. Genome-wide identification of tebufenozide resistant genes in the smaller tea tortrix, *Adoxophyes honmai* (Lepidoptera: Tortricidae). Sci Rep 9:4203.

41. Suzuki MG, Funaguma S, Kanda T, Tamura T, Shimada T. 2003. Analysis of the biological functions of a doublesex homologue in *Bombyx mori*. Dev Genes Evol 213:345–354.

42. Suzuki MG, Funaguma S, Kanda T, Tamura T, Shimada T. 2005. Role of the male BmDSX protein in the sexual differentiation of *Bombyx mori*. Evol Dev 7:58–68.

43. Clough E, Jimenez E, Kim YA, Whitworth C, Neville MC, Hempel LU, Pavlou HJ, Chen ZX, Sturgill D, Dale RK, Smith HE, Przytycka TM, Goodwin SF, Doren MV, Oliver B. 2014. Sex-and tissue-specific functions of *Drosophila* doublesex transcription factor target genes. Dev Cell 31:761–773.

44. Cools R. 2006. Dopaminergic modulation of cognitive function-implications for L-DOPA treatment in Parkinson’s disease. Neurosci Biobehav Rev 30:1–23.

45. Vavricka C, Han Q, Huang Y, Erickson SM, Harich K, Christensen BM, Li J. 2011. From L-dopa to dihydroxyphenylacetaldehyde: a toxic biochemical pathway plays a vital physiological function in insects. PLoS One 6:e16124.

46. Kinjoh T, Kaneko Y, Itoyama K, Mita K, Hiruma K, Shinoda T. 2007. Control of juvenile hormone biosynthesis in *Bombyx mori:* cloning of the enzymes in the mevalonate pathway and assessment of their developmental expression in the corpora allata. Insect Biochem Mol Biol 37:808–818.

47. Saiki S, Sasazawa Y, Fujimaki M, Kamagata K, Kaga N, Taka H, Li Y, Souma S, Hatano T, Imamichi Y, Furuya N, Mori A, Oji Y, Ueno S, Nojiri S, Miura Y, Ueno T, Funayama M, Aoki S, Hattori N. 2019. A metabolic profile of polyamines in parkinson disease: A promising biomarker. Ann Neurol 86:251–263.

48. Nunomura A, Moreira PI, Lee HG, Zhu X, Castellani RJ, Smit MA, Perry G. 2007. Neuronal death and survival under oxidative stress in Alzheimer and Parkinson diseases. CNS Neurol Disord Drug Targets 6:411–423.

49. Hoshino M, Nakanishi K, Nakai M, Kunimi Y. 2008. Gross morphology and histopathology of male-killing strain larvae in the oriental tea tortrix *Homona magnanima* (Lepidoptera: Tortricidae). Appl Entomol Zool 43:119–125.

50. Charlat S, Davies N, Roderick GK, Hurst GDD. 2007. Disrupting the timing of *Wolbachia-induced* male-killing. Biol Lett 3:154–156.

51. Sahara K, Yoshido A, Traut W. 2012. Sex chromosome evolution in moths and butterflies. Chromosome Res 20:83–94.

52. Kageyama D, Ohno M, Sasaki T, Yoshido A, Konagaya T, Jouraku A, Kuwazaki S, Kanamori H, Katayose Y, Narita S, Miyata M, Riegler M, Sahara K. 2017. Feminizing *Wolbachia* endosymbiont disrupts maternal sex chromosome inheritance in a butterfly species. Evol Lett 1:232–244.

53. Kageyama D, Hoshizaki S, Ishikawa Y. 1998. Female-biased sex ratio in the Asian corn borer, *Ostrinia furnacalis:* evidence for the occurrence of feminizing bacteria in an insect. Heredity 81:311–316.

54. Arai H, Ishitsubo Y, Nakai M, Inoue MN. 2022. Mass-Rearing and Molecular Studies in Tortricidae Pest Insects. J Vis Exp.

55. Grabherr MG, Haas BJ, Yassour M, Levin JZ, Thompson DA, Amit I, Adiconis X, Fan L, Raychowdhury R, Zeng Q, Chen Z, Mauceli E, Hacohen N, Gnirke A, Rhind N, Palma F, Birren BW, Nusbaum C, Lindblad-Toh K, Friedman N, Regev A. 2011. Full-length transcriptome assembly from RNA-Seq data without a reference genome. Nat Biotechnol 29:644–652.

56. Bray NL, Pimente, H, Melsted P, Pachter L. 2016. Near-optimal probabilistic RNA-seq quantification. Nat Biotechnol 34:525–527.

57. Conesa A, Götz S, García-Gómez JM, Teról J, Talon M, Robles M. 2005. Blast2GO: a universal tool for annotation, visualization and analysis in functional genomics research. Bioinform 21:3674–3676.

58. Kawamoto M, Jouraku A, Toyoda A, Yokoi K, Minakuchi Y, Katsuma S, Fujiyama A, Kiuchia T, Yamamoto K, Shimada T. 2019. High-quality genome assembly of the silkworm, *Bombyx mori*. Insect Biochem Mol Biol 107:53–62.

59. Gu L, Walters JR, Knipple DC. 2017. Conserved patterns of sex chromosome dosage compensation in the Lepidoptera (WZ/ZZ): insights from a moth neo-Z chromosome. Genome Biol Evol 9:802–816.

60. Staley K, Blaschke AJ, Chun J. 1997. Apoptotic DNA fragmentation is detected by a semi-quantitative ligation-mediated PCR of blunt DNA ends. Cell Death Differ 4:66–75.

61. Arai H, Ishitsubo Y, Nakai M, Inoue MN. 2022. A simple method to disperse eggs from lepidopteran scalelike egg masses and to observe embryogenesis. Entomol Sci 25:e12497.

